# *In situ* structures of periplasmic flagella in *Borrelia* reveals conformational changes essential for flagellar rotation

**DOI:** 10.1101/553289

**Authors:** Yunjie Chang, Kihwan Moon, Xiaowei Zhao, J. Norris Steven, Md A. Motaleb, Jun Liu

## Abstract

The bacterial flagellar motor is a molecular machine that rotates the flagellar filament at high speed. Torque is generated by the stator-rotor interaction coupled to an ion flux through the torque-generating stator. Here, we employed cryo-electron tomography to visualize the intact flagellar motor in the Lyme disease spirochete *Borrelia burgdorferi*. By analysis of the motor structures of wild-type and stator mutants, we localize the torque-generating units precisely and determine three-dimensional structure of the stator and its interactions with the rotor. Our study shows that the cytoplasmic domains of the stator are packed regularly around the circumference of the flagellar C-ring. The stator-rotor interaction induces a profound conformational change in the C-ring. Analysis of the motors of a less motile *motB-*D24E mutant and a non-motile *motB-*D24N mutant, in which the proton translocation is reduced and blocked, respectively, provides evidence that the conformational change of the C-ring is essential for flagellar rotation.

## INTRODUCTION

The bacterial flagellum is the core organelle of swimming motility, and this motility is often important for the virulence of bacterial pathogens. The flagellum is composed of at least 25 different proteins that form a rotary motor, a hook, and a long filament ^*1-3*^. The rotation of the motor is driven by ion-motive forces: the flagellar motors of most bacteria (including the model organism *E. coli*) use a proton-motive force (PMF) ^*4*^, whereas the polar flagella in *Vibrio* use a sodium-motive force (SMF) ^*5*^. Torque is generated by ion flow through the membrane-spanning, torque-generating units known as the stators ^*2, 6, 7*^.

In many bacteria, rotor proteins FliG, FliM and FliN constitute the C-ring, which transmits the torque from the stator to rotate the MS-ring, rod, hook, and filament. In H^+-^driven motors, the stators are formed by the MotA and MotB proteins in a MotA_4_MotB_2_ transmembrane complex, which contains two proton channels ^*8-10*^. Each MotA has four transmembrane segments (TM1-TM4) and a large cytoplasmic loop, which contains several conserved charged residues that are critical for interactions with the C-ring ^*11*^. MotB consists of one transmembrane segment and a large periplasmic domain that contains a putative peptidoglycan-binding (PGB) motif ^*12*^. The stator complex can be inactivated when its ion channels are plugged by a short, amphipathic helix, called the lid, which prevents protons flowing into the cytoplasm until the rotor is engaged with the C-ring ^*13*^.

Once a stator complex incorporates into a motor, a substantial conformational change occurs in the periplasmic domain of MotB to allow attachment to the PG layer and to open the proton channel ^*14-16*^. A conserved aspartyl residue (Asp32 in *Escherichia coli* MotB, Asp33 in *Salmonella enterica* MotB) in the transmembrane segment of MotB is predicted to be the proton-binding site ^*17*^. It is thought that proton binding/dissociation at this residue triggers conformational changes in the cytoplasmic domain of MotA to produce a power-stroke on the C-ring that drives flagellar rotation ^*18*^.

Fluorescence recovery after photobleaching (FRAP) analysis in *E. coli* showed that stators associated with the C-ring constantly exchange with stators in the membrane pool ^*19*^. The observation of stepwise increases and decreases between discrete speeds in a single motor provides further evidence for the dynamic nature and variable number of the torque-generating units in the *E. coli* motor ^*20-22*^. Ion flow is also important for the assembly of the torque-generating units in *E. coli* and *Vibrio alginolyticus* ^*23, 24*^, as its disruption leads to reversible stator diffusion away from the motor. Furthermore, the charged residues on the cytoplasmic loop of MotA are required for stator assembly in the flagellar motor of *S. enterica* ^*25*^. The C-terminal domain of FliG (FliG_C_) is also critical for the proper assembly of PomA/PomB stator complex in *V. alginolyticus* flagellar motor; several point mutations at the FliG_C_ completely abolish both motility and polar localization of the stator without affecting flagellation ^*26*^. All of these phenotypes are consistent with the dynamic nature of the torque-generating units.

The initial images of the torque-generating units were shown by electron micrographs of freeze-fractured membrane, which revealed between 10 and 12 stud-like particles in *E. coli*, and between 14 and 16 particles in *Streptococcus* and *Aquaspirillum serpens*, packing around the flagellar base ^*27, 28*^. A 3-D structure of a stator was reconstructed from a purified PomA/PomB complex ^*29*^. The reconstructed map showed two arm-like periplasmic domains and a large cytoplasmic domain. However, the isolated stator without its context is insufficient to determine how the torque-generating units are arranged in an intact flagellar motor.

Recently, cryo-electron tomography (cryo-ET) has provided a new approach to study the intact flagellar motor in bacteria ^*30-34*^. However, a detailed visualization of the torque-generating units remains difficult in many bacterial species, such as *E. coli* or *S. enterica*, partly because the stators are highly dynamic and also because wild type cells are too large for high resolution cryo-ET imaging. In contrast, stable putative torque-generating units have been described in some flagellar motors, including the spirochetal periplasmic motors ^*30-36*^. Presumably, the stators in these motors either do not exchange or exchange occurs without disrupting the overall arrangement of the torque generator units. However, the relatively low resolution of previous images has posed challenges to identifying the *in situ* torque-generating stators and to reveal the precise interaction between stators and other motor components.

Spirochetes represent one of the major bacterial phyla. They have an unusual cell morphology and form of motility. The flagella are enclosed between the outer membrane and the PG layer, and are thus called periplasmic flagella (PFs). The flagellar motors are found at both cell poles and rotate coordinately to enable the cell to run, pause or flex ^*37*^. *Borrelia burgdorferi*, one of the agents of Lyme disease, is a highly motile and invasive spirochete pathogen. There are 7-12 PFs anchored at each cell pole. The rotation of PFs causes the cell to form a flat-wave shape to efficiently bore its way through tissue and viscous media ^*37-40*^.

Although the *B. burgdorferi* motor differs in some aspects from the *E. coli* motor (e.g. the presence of a prominent “collar” structure), genome sequence analyses as well as in situ structural analyses suggest that the major flagellar components, including the MS-ring, the C-ring, the rod, the export apparatus, the hook, and the filament are remarkably similar to those of other bacterial species ^*36*^. The *B. burgdorferi* motor has been dissected by construction of many mutants, either by transposon mutagenesis or in-frame deletion. Our study of *fliE, flgB, flgC, flhO* and *flgG* deletion mutants yielded insights into rod assembly ^*41*^. Studies of *fliI* and *fliH* mutants provided unprecedented structural information about the export apparatus and their role in the assembly of PFs ^*42, 43*^. Furthermore, the membrane protein FliL was identified as an important player that controls the organization of the PFs ^*44*^. More recently, the collar (a spirochete-specific feature) was determined to play an important role in recruiting and stabilizing the stators ^*45, 46*^. Thus, *B. burgdorferi* has emerged as a tractable model organism to study the structure and function of bacterial flagella at molecular resolution ^*36*^.

In this study, we focused primarily on determining the assembly and interaction of the torque-generating stator with other motor components of *B. burgdorferi*. By comparative analysis of the motor structures from wild-type, mutants, complemented mutant strains, we localized the torque-generating units in the spirochetal flagellar motor. Importantly, detailed analysis of the stator-rotor interaction revealed a novel conformational change that is required to transmit torque from the stator to the C-ring.

## RESULTS

### The MotA/MotB complex is the torque-generating unit in ***B. burgdorferi***

The MotA/MotB complex is the torque generator in PMF-driven flagellar motors, as knockout mutations in *motA* or *motB* of *E. coli* eliminate motility without affecting flagellar assembly ^*21*^. The *motA* and *motB* genes are located in the middle of the *flgB* operon in *B. burgdorferi* (Figure 1A). Our recent study reported inactivation of *motB* in *B. burgdorferi* and its complementation *in trans* ^*47*^. The δ*motB* cells were completely non-motile and had an irregular, rod-shaped morphology, very different from the wave-like, highly motile wild type (WT) and complemented cells (*motB*^com^) (Figure 1C, Figure 1-figure supplement 1). The Δ*motB* cells possess paralyzed flagella, suggesting that the Δ*motB* mutant lacks the torque-generating unit (Figure 1D). The Δ*motA* mutant exhibited similar motility and morphology phenotypes (data not shown), consistent with the notion that MotA and MotB form a complex for torque generation ^*48*^.

**Figure 1.**
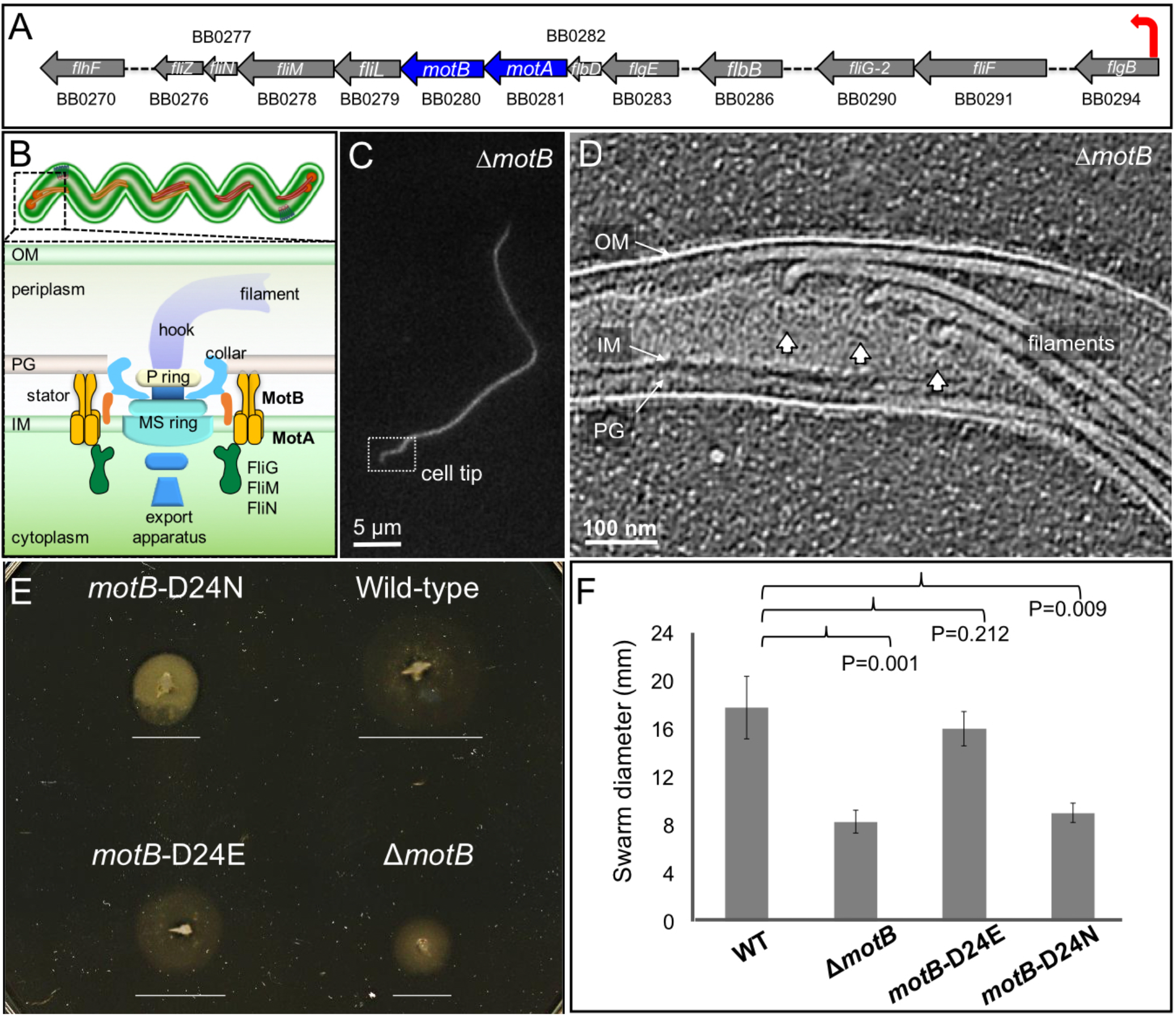
Overview of flagella organization in *Borrelia burgdorferi* and the motility phenotypes of of WT, Δ*motB*, and point mutants of *motB*. (A) Schematic of the *flgB* flagellar operon map of *B. burgdorferi*. Red arrow indicates the direction of transcription. The *motA* and *motB* genes are shown as blue arrows. (B) Schematic models of a periplasmic flagellum, including the motor. (C) A typical dark-field microscopy image of a Δ*motB* cell. (D) A section from a typical tomogram of a Δ*motB* cell tip shows multiple flagellar motors and filaments *in situ*. (E) Swarm plate assay of Δ*motB, motB*-D24E, *motB*-D24N, and WT cells. (F) Averages ± standard deviations of swarm diameters from WT, Δ*motB, motB*-D24E and *motB*-D24N strains. A paired Student’s *t* test was used to determine a *P* value. A P<0.05 between strains is considered significant.

In *B. burgdorferi* MotB, Asp-24 is the homologous residue to the highly conserved aspartate (Figure 1-figure supplement 2) that is essential for proton translocation in *E. coli* and *S. enterica* ^*17, 49*^. To examine the role of proton transport in *Borrelia* stator assembly and activity, we introduced point mutations to generate *motB*-D24E and *motB*-D24N. Dark-field microscopy and swarm plate motility assays indicated that *motB*-D24E mutant cells are less motile than the WT cells, and *motB*-D24N cells are completely non-motile (Figure1E, F). The result obtained with *motB*-D24E is consistent with the reduced motility observed with the D32E substitution in *E. coli* MotB ^*17*^ and the D33E substitution in *S. enterica* ^*49*^. The *B. burgdorferi motB*-D24N phenotype is also identical to that of the D32N substitution in *E. coli* MotB ^*17, 50*^. Based on these and prior results ^*47*^, we conclude that the MotA/MotB complex in *B. burgdorferi* is the torque-generating stator and that Asp-24 in *B. burgdorferi* MotB is essential for proton translocation and motility.

### Characterization and localization of the torque-generating units in intact ***B. burgdorferi* motor**

To image the torque-generating units and their interactions with other flagellar proteins *in situ* at the molecular level, we generated asymmetric reconstructions of the flagellar motor from Δ*motB* and Δ*motA* strains (Figure 2 and Figure 2-figure supplement 1). The averaged structures of the motors in the Δ*motB* and Δ*motA* mutants exhibited many of the same features as the WT motor, such as the export apparatus, the C-ring, the MS-ring, the rod, the P-ring, and the spirochete-specific periplasmic collar. However, compared to the WT motor structure, the Δ*motA* and Δ*motB* mutants lack large transmembrane densities peripheral to the C-ring (indicated by arrows in Figure 2, Video 1). Complementation of *motB* restored the missing densities in the Δ*motB* mutant (Figure 2-figure supplement 1) and also restored motility and wild-type cell morphology (Figure 1-figure supplement 1). These results are consistent with the peripheral densities in the WT motor being comprised of MotA/MotB complexes, as has been found with other bacteria. In each WT motor, sixteen of these densities are symmetrically distributed around the C-ring (Figure 2E). They form a stator ring with a diameter of ∼80 nm, which is significantly larger than the C-ring with a diameter of ∼57 nm (Figure 2E, Video 1). The WT flagellar C-ring has 46-fold symmetry (Figure 3E and Figure 3-figure supplement 1), while the surrounding stators have a 16-fold symmetry. Therefore, there is a symmetry mismatch between the C-ring and its surrounding stators.

**Figure 2.**
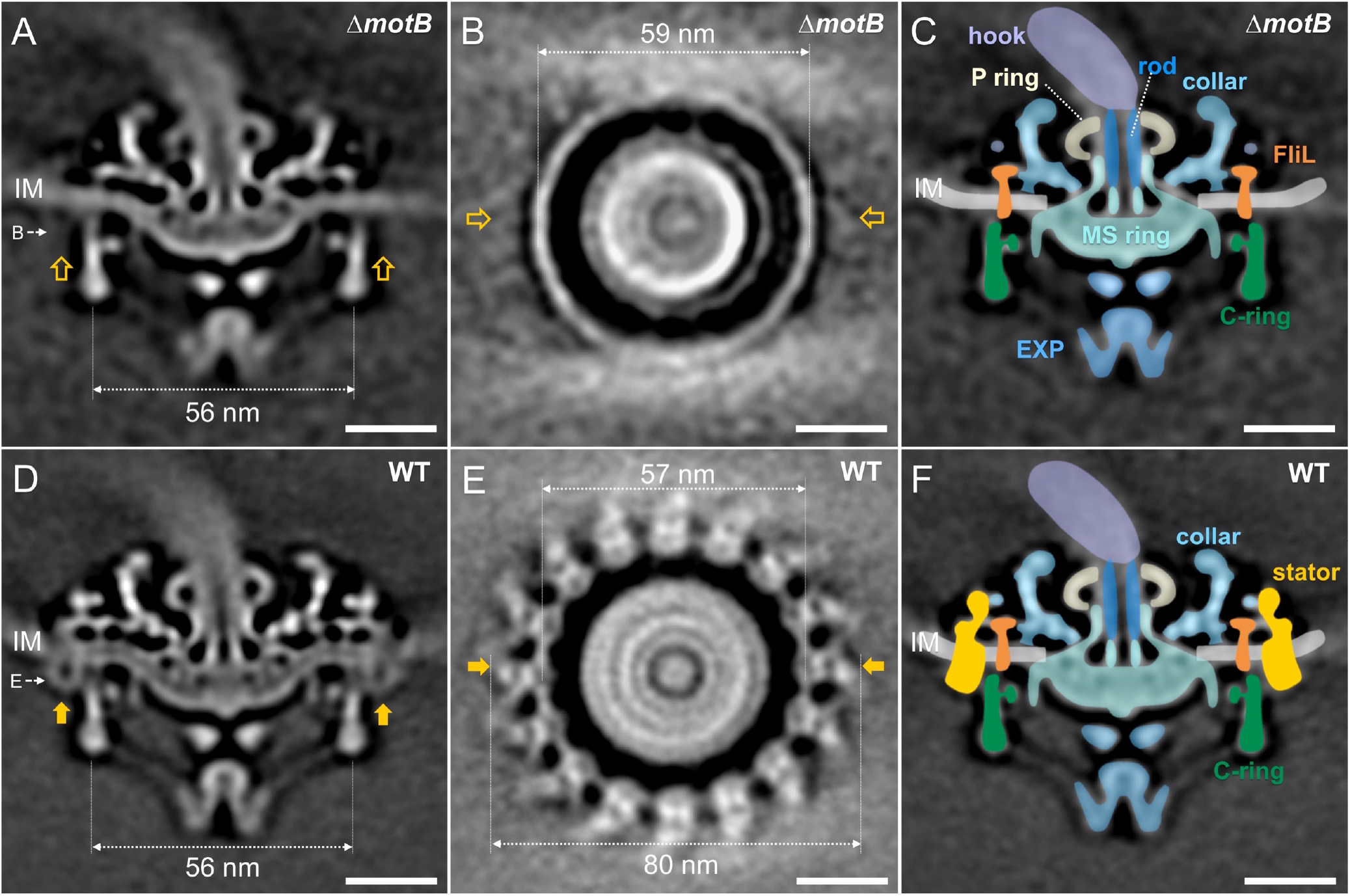
Asymmetric reconstructions of Δ*motB* and WT motors in *B. burgdorferi*. (A) A central section of the flagellar motor structure from a Δ*motB* mutant. The diameter of the bottom of the C-ring is 56 nm. The missing densities compared to the WT flagellar motor are indicated by empty arrows. (B) A cross-section at the top of the C-ring from the Δ*motB* flagellar motor structure. The diameter of the top of the C-ring is 59 nm. (C) A cartoon model highlights key components in the Δ*motB* flagellar motor: C-ring (green), export apparatus (EXP), MS-ring (blue-green) embedded in the inner membrane (IM), FliL (coral), collar (light blue), P-ring (gray), and rod (blue). (D) A central section of the flagellar motor structure from WT. The diameter of the bottom of the C-ring is 56 nm. The extra densities compared to Δ*motB* flagellar motor structure are indicated by solid orange arrows. (E) A cross-section at the top of the C-ring from the WT flagellar motor structure. The diameter of the top of C-ring is 57 nm. Note that there are sixteen stator densities associated with the C-ring. The diameter of the stator ring is 80 nm. (F) A cartoon model highlights key flagellar components in the WT flagellar motor: C-ring, MS-ring, FliL, collar, and stators embedded in the IM. Scale bar = 20 nm.

**Figure 3.**
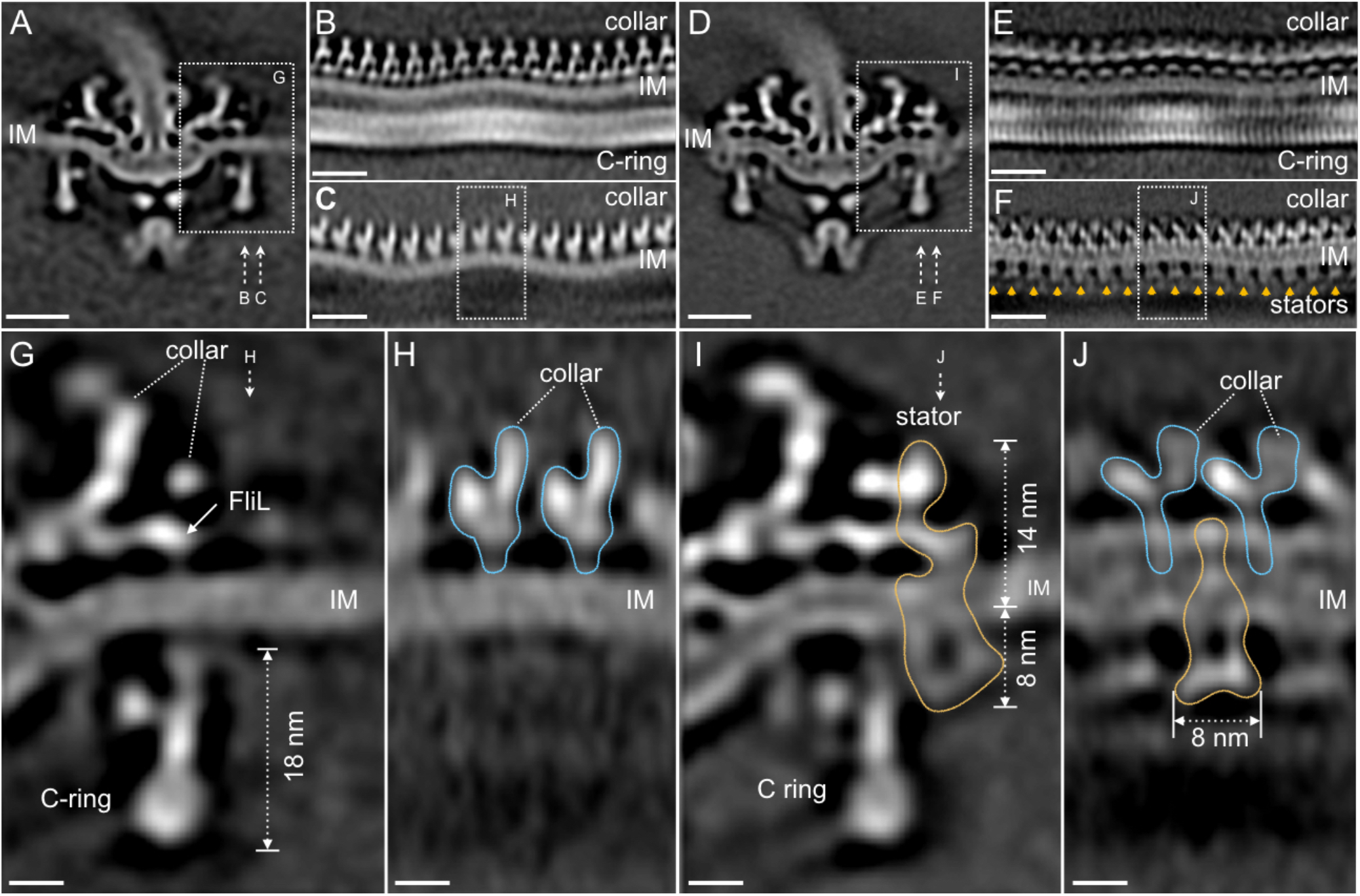
Visualization of the stators and their interactions with the periplasmic components and the C-ring. (A) A central section of the Δ*motB* motor structure. (B, C) Two sections from an unrolled map of the Δ*motB* motor showing the curvature of the inner membrane (IM), the intrinsic flexibility of the collar and the C-ring, respectively. (D) A central section of the WT motor structure. (E) One section from the unrolled map of the WT motor showing the symmetry mismatch between the C-ring and the collar. (F) Another section from the unrolled map of the WT motor showing sixteen stators (indicated by orange arrows) embedded in the IM. (G) A central section from a refined structure of the Δ*motB* motor showing the C-ring, collar and FliL embedded in the IM. (H) A perpendicular section showing the collar (blue line highlighted) on top of the IM. (I) A central section from a refined structure of the WT motor showing the C-ring, and the stator (outlined by a gold line). (J) A perpendicular section showing the stator (outlined by a gold line) inserted between two subunits of the collar (highlighted by blue lines).

### The stator-rotor interaction induces large conformational changes in the C-ring

In the presence of the stators, the top portion of the C-ring undergoes considerable changes. Specifically, there is ∼2 nm shrinkage in diameter (59 nm vs. 57 nm) from the Δ*motB* mutant motor to the WT motor (Figs. 2B, 2E). As the entire motor is embedded in the highly curved membrane of the spirochete, the C-ring, the peripheral part of the collar and its underlying membrane appear to be variable in height in both *ΔmotB* and WT motors (Figure 3B, C). The sixteen stators in the WT motor also display considerable variation in height to be embedded in the cytoplasmic membrane and inserted between the “collar” (Figure 3H).

To obtain a high-resolution structure of the stator and its interaction with the C-ring, the sixteen stator units in each averaged motor structure were rotationally aligned and classified. Comparative analysis of the class averages derived from Δ*motB* and WT motors enabled us to determine the *in situ* structure of an individual stator. The stator is composed of a periplasmic domain and a cytoplasmic domain. The periplasmic domain is ∼14 nm in height, and it directly interacts with the collar and FliL (Figure 3I), which have previously been identified as periplasmic structures ^*44*^. The interactions are believed to play critical roles in stabilizing the stator, as the lack of FliL or collar proteins has a profound impact on stator assembly and motility ^*44-46*^. Notably, the collar-FliL assembly also exhibits conformational changes due to the presence or absence of stators (Figure 3H, J), further supporting the strong protein-protein interactions among the stator, FliL, and collar (Figure 4A, B).

**Figure 4.**
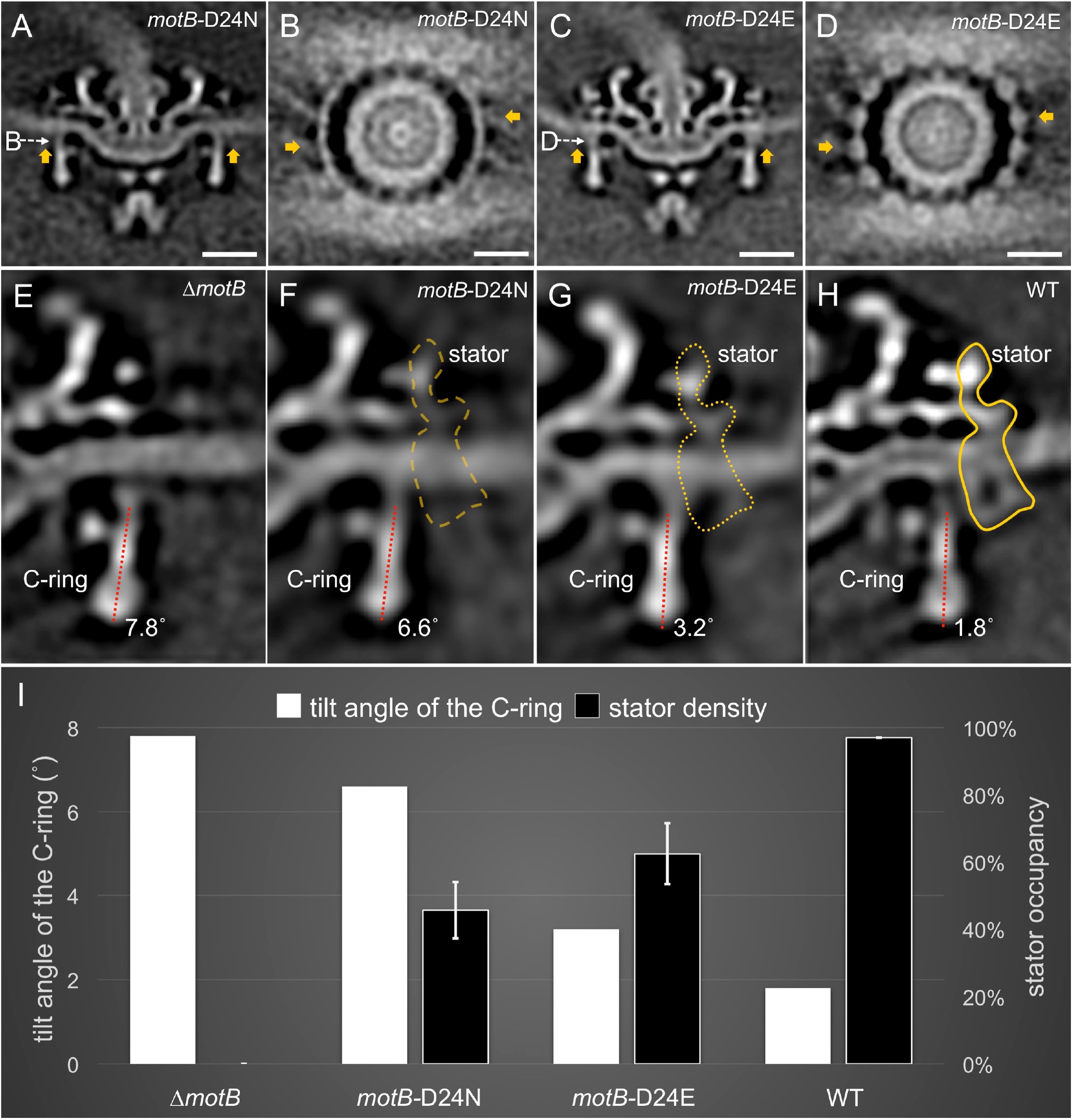
Stator binding and proton flux are required to induce the conformational changes of the C-ring. (A) A central section from the asymmetric reconstruction of the *motB*-D24N motor. (B) A cross-section from the asymmetric reconstruction of the *motB*-D24N motor at the interface between the C-ring and the stators. (C) A central section from the asymmetric reconstruction of the *motB*-D24E motor. (D) A cross-section from the asymmetric reconstruction of the *motB*-D24E motor at the interface between the C-ring and the stators. Two stators are indicated by arrows in (A-D). For a better comparison, we showed the central sections from the refined Δ*motB* motor structure (E), the refined *motB*-D24N motor structure (F), the refined *motB*-D24E motor structure (G) and the refined WT motor structure (H), respectively. Note that the stator densities in the *motB*-D24E and *motB*-D24N mutants are considerably weaker than that in the WT motor, while there is no stator density in the Δ*motB* mutant (E). The tilt angle of the C-ring away from the rotation axis is about 7.8° in the Δ*motB* motor (E), while those in the *motB*-D24N, *motB*-D24E and WT motors are 6.6°, 3.2°, 1.8°, respectively. (I) 3D classification based on the C-ring density and stator density reveals various conformations of the C-ring and different occupancy of the stator in four strains: WT, *motB*-D24E, *motB*-D24N, and Δ*motB*.

The cytoplasmic domain of the stator is ∼8 nm in diameter, which is comparable to the diameter observed in the freeze-fractured membrane ^*27, 28, 51*^. They are adjacent to the top portion of the C-ring (Figure 3I). Importantly, the interaction between the stator and the C-ring induced a noticeable conformational change of the C-ring compared to that in Δ*motB* mutant motor (Figure 3G). Specifically, the top of the C-ring is tilted toward the MS-ring by ∼6° in the presence of the stator (Figure 3I). This ‘tilt’ corresponds with the ∼ 2 nm reduction in the diameter of the top portion of the C-ring (Figure 2B, E) observed in WT relative to Δ*motB* mutant motors.

### The conformational changes of the C-ring are directly linked to higher torque and faster motility

To gain a better understanding of the conformational changes in the C-ring that accompany assembly of the torque-generating units, we examined the motor structures in the less-motile *motB*-D24E mutant and the non-motile *motB*-D24N mutant and compared them with the motor structures from WT and the Δ*motB* mutant (Figure 4). Both the *motB*-D24E and *motB*-D24N mutants have stators assembled in the motor (Figure 4A-D), but the stator densities are significantly weaker than that in WT cells (Figure 2D), suggesting that the stators are variable in their location or occupancy. Statistical analysis of the stators showed that the stator occupancy is 97.0% in WT organisms, suggesting that there are ∼16 stators in each WT motor. In the *motB*-D24E motor, the corresponding stator density was ∼62.5%, suggesting that there are ∼10 stators in average. In *motB*-D24N, there were an estimated ∼7 stators per motor in average (Figure 4 and Figure 4-figure supplement 1). As these two residue substitutions are known to reduce or block (respectively) proton flux in the stator channel, reduced stator occupancy in these mutants is consistent with the idea that PMF-mediated proton translocation affects MotB conformation and protein interactions involved in stator assembly and stability ^*23, 24*^.

The motility phenotypes of cells expressing two different mutant motors are very different. The *motB*-D24E cells are less motile than the WT, whereas *motB*-D24N cells are non-motile. The C-ring undergoes a considerable change at the stator binding positions in the *motB*-D24E cells (Figure 4B), as in the WT (Figure 2E). However, there is no visible change observed in the *motB-*D24N cells (Figure 4D). 3D classification of the C-ring densities shows that the averaged value of the C-ring tilt angle in *motB-*D24E and *motB-*D24N motors is 3.2° and 6.6°, respectively (Figure 4F, G and Figure 4-figure supplement 2). In contrast, the C-ring in Δ*motB* motors has an oblique angle of ∼7.8°, while the C-ring in WT is tilted at ∼1.8° (Figure 4E, H and Figure 4-figure supplement 2). These results indicate that the conformational change of the C-ring is related to the stator quantity and proton flux transmitting through the stator channel. Consequently, increased number of stator units and higher torque resulted in larger conformational change of the C-ring.

### Model of stator-rotor interactions in ***B. burgdorferi***

To model the stator-rotor interactions and the conformational change of the C-ring in detail, we first built pseudo-atomic structures of the C-ring in the Δ*motB* mutant, as it is structurally similar to the C-ring of *S. enterica*. The FliN tetramer was placed into the bulge at the bottom of the C-ring as proposed in *E. coli* ^*52*^. The FliM_M_-FliG_MC_ complex (PDB: 4FHR) ^*53*^ and the N-terminal domain of FliG (PDB 3HJL) ^*54*^ were docked onto the top portion of the C-ring. The densities were well fitted with 46 FliG proteins organized in a ring in our EM map (Figure 5, Vidoe 2). Noticeably, FliG proteins are relatively far away from the periphery of the MS-ring and the C-ring appears to be disengaged from the MS-ring.

**Figure 5.**
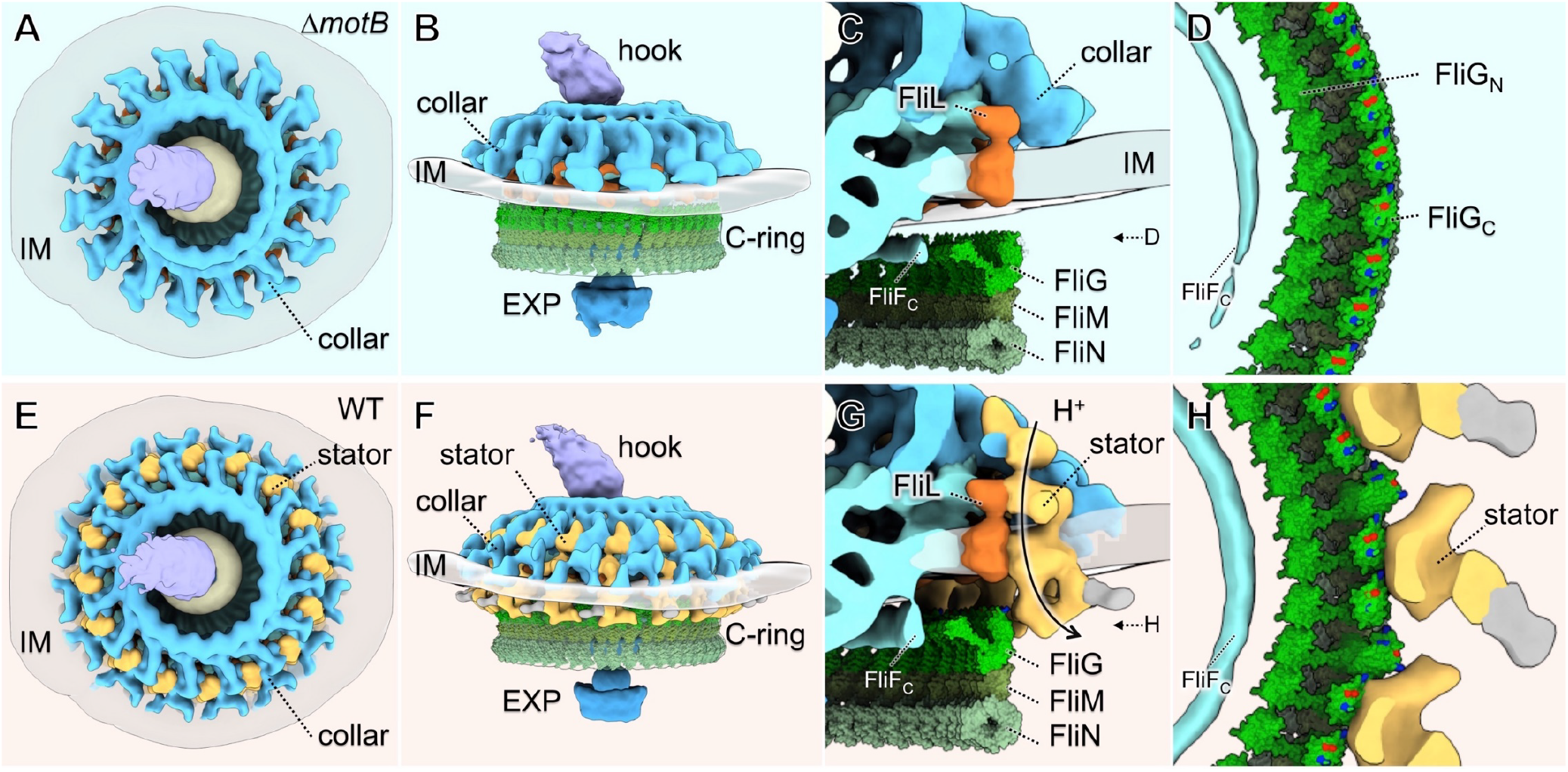
Molecular architecture of the stator-rotor interactions. (A, B) A top and a side view showing the surface rendering of the Δ*motB* motor, respectively. (C) A side view shows major flagellar components: the cytoplasmic domain of FliF (FliF_C_), FliL, FliG, FliM, FliN and the collar around the inner membrane (IM). (D) A top view of the interface between the C-ring and the MS-ring. FliF_C_ of the MS-ring is adjacent to the FliG_N_ of the C-ring. (E, F) A top and a side view showing the surface rendering of the WT motor. (G) A side view of the interface between the stator and the C-ring. The interaction powered by proton flux induce conformational change of the C-ring, which appears to engage with the MS-ring through interactions between FliG_N_ and FliF_C_. (H) A top view of the interface between the C-ring and the stators. The charged residues in FilG_C_ are shown in blue (positive electrostatic potential) or red (negative electrostatic potential).

To build the C-ring model in the WT motor, we rotated the FliG/FliN/FliM complex as a rigid body toward the MS-ring for about ∼6° (Figure 5, Video 2). The additional shifts of the FliG proteins enables the C-ring to engage with the MS-ring at its periphery. As the result, the N-terminal domain of FliG interacts with the periphery of the MS-ring and the C-terminal domain of FliG interacts with the stator. Specifically, two charged residues in the C-terminal domain of FliG are adjacent to the cytoplasmic portion of the stator, presumably interacting with the charged residues of MotA (Figure 5). Therefore, our *in situ* structures are consistent with the previous findings in *E. coli* motor that these charged residues in FliG and the cytoplasmic loop of MotA are important for torque generation ^*11, 55, 56*^.

## DISCUSSION

The bacterial flagellum is one of the most fascinating prokaryotic motility organelles. It has been studied extensively in the model systems *E. coli* and *S. enterica* Typhimurium for several decades. However, our understanding of flagellar assembly and rotation remains incomplete, partly because the torque-generating units in Enterobacteriaceae are highly dynamic and limited structural information about the torque-generating units and their interactions with the rotor are available. Recent studies provide clear evidence that flagella from different bacterial species have evolved considerable variation in flagellar structure as well as in flagellar number and placement to adapt to the specific environments that bacteria encounter. Spirochetes are unique in their evolution of periplasmic flagella, resulting in a form of locomotion effective in viscous environments such as dense mud and host tissue. Their flagellar motors are not only significantly larger than those found in *E. coli* and *S.* Typhimurium, but also possess a unique periplasmic collar structure. In this study, the relative stability and high occupancy rate of the *B. burgdorferi* stator together with the availability of key mutants permitted the use of cryo-ET analysis to reveal the stator structure and its interactions with the rotor at a high resolution. This approach will likely contribute to our understanding the mechanisms underlying flagellar motor assembly and rotation.

### The unique structure of periplasmic flagella is critical to their function

The periplasmic collar constitutes a large, turbine-like complex in the flagellar motor of spirochetes. This structure plays an important role in the assembly of periplasmic flagella and hence in determining cell morphology and motility ^*45, 46*^. In contrast to the highly dynamic stators within the *E. coli* flagellar motor, sixteen torque-generating units appears to be stably assembled around the collar in *B. burgdorferi* (Figure 2 and Figure 3). Therefore, the collar is likely to form a scaffold for assembling and stabilizing the torque-generating stator units. In *B. burgdorferi*, FliL also forms additional feature located between the stator and the collar in periplasm ^*44*^. Deletion of *fliL* resulted in cells with defects in motility ^*44*^ and a flagellar motor with fewer stator units (Figure 4-figure supplement 1, Figure 4-figure supplement 4) compared to that in the WT motor, suggesting that FliL also plays an important role in stator assembly in *B. burgdorferi*.

### Impact of PMF on stator assembly and C-ring conformation in periplasmic flagella

PMF is not only essential for flagellar rotation but also critical for assembly of the stators around the motor in *E. coli* and other model systems. By altering the putative proton channel in *B. burgdorferi*, we found that the average number of stators is decreased significantly in the *motB*-D24E (65% occupancy] and *motB*-D24N (45% occupancy) motors (Figure 4). However, even in the non-motile *motB*-D24N cells, some stators remain associated with the motor. This finding is different from the observations in *E. coli* and other model systems, in which stators were found to dissociate from both Na^+^- and H^+^-driven motors when the IMF was disrupted ^*23, 24*^. The periplasmic collar in spirochetes may be the key reason underlying the difference, as deletion of genes that encode the proteins of the collar disrupt the assembly of the stators ^*45, 46*^.

It has been proposed that proton flow through the motor triggers conformational changes in the stator that generate a power stroke to the C-ring ^*18*^. However, it has been difficult to directly observe the conformational changes, partly because of the dynamic nature of the stators and their interactions with the C-ring. To visualize the conformational changes in detail, we took advantage of several unique aspects in *B. burgdorferi* flagellar system. 1) The large periplasmic collar and FliL help to recruit sixteen stators to each motor. 2) A high torque is required to drive the rotation of periplasmic flagella and of the entire body. 3) Multiple motors located in at the poles of skinny cells enable high-resolution cryoET imaging. 4) Recent advances in the genetic tools available for *B. burgdorferi* enable us to make specific mutants in different flagellar components. As a result, we were able to observe a large conformational change in the *B. burgdorferi* flagellar C-ring due to stator assembly, proton transport, and resulting rotation. Importantly, stator binding alone is not sufficient to induce the C-ring conformational changes, because stators associated with the *motB*-D24N motor are not able to interact effectively with the C-ring and drive flagellar rotation. As proton conduction is blocked in the *motB*-D24N motor, we believe that proton conduction is not only essential for the generation of the power stroke, but also induces conformational changes in the C-ring needed for rotational activity. Although we did not see the conformation changes in the stator as proposed previously ^*18*^, we observed its impact as indicated by the C-ring conformational change, which is presumably required for flagellar rotation.

### A model of stator-rotor interactions in ***B. burgdorferi***

Our *in situ* structural analysis of *B. burgdorferi* flagellar motors enables us to propose a model for stator assembly and stator-rotor interactions. Before assembly of the stators, the *B. burgdorferi* flagellar motor is composed of the MS-ring, the rod, the export apparatus, the collar with 16-fold symmetry, and the C-ring with 46-fold symmetry (Figure 6). The collar and the associated FliL protein provide 16 well-defined locations for the recruitment of stator complexes, which assemble around the collar and the C-ring. Stator complexes with decreased or blocked proton conduction only partially occupy the 16 possible locations (Figure 6). There is little conformational change in the C-ring at the stator-rotor interface. With increases in proton conduction, as in the wild-type Asp-24 stator, all 16 positions become occupied. The deformation of the C-ring - the tipping shown in Figure 4 - increases with increased proton-conduction capability and is highest in C-rings surrounded by wild-type stators (Figure 6). This tipping may therefore represent conformational changes in the C-ring that are consequences of torque-generation.

**Figure 6.**
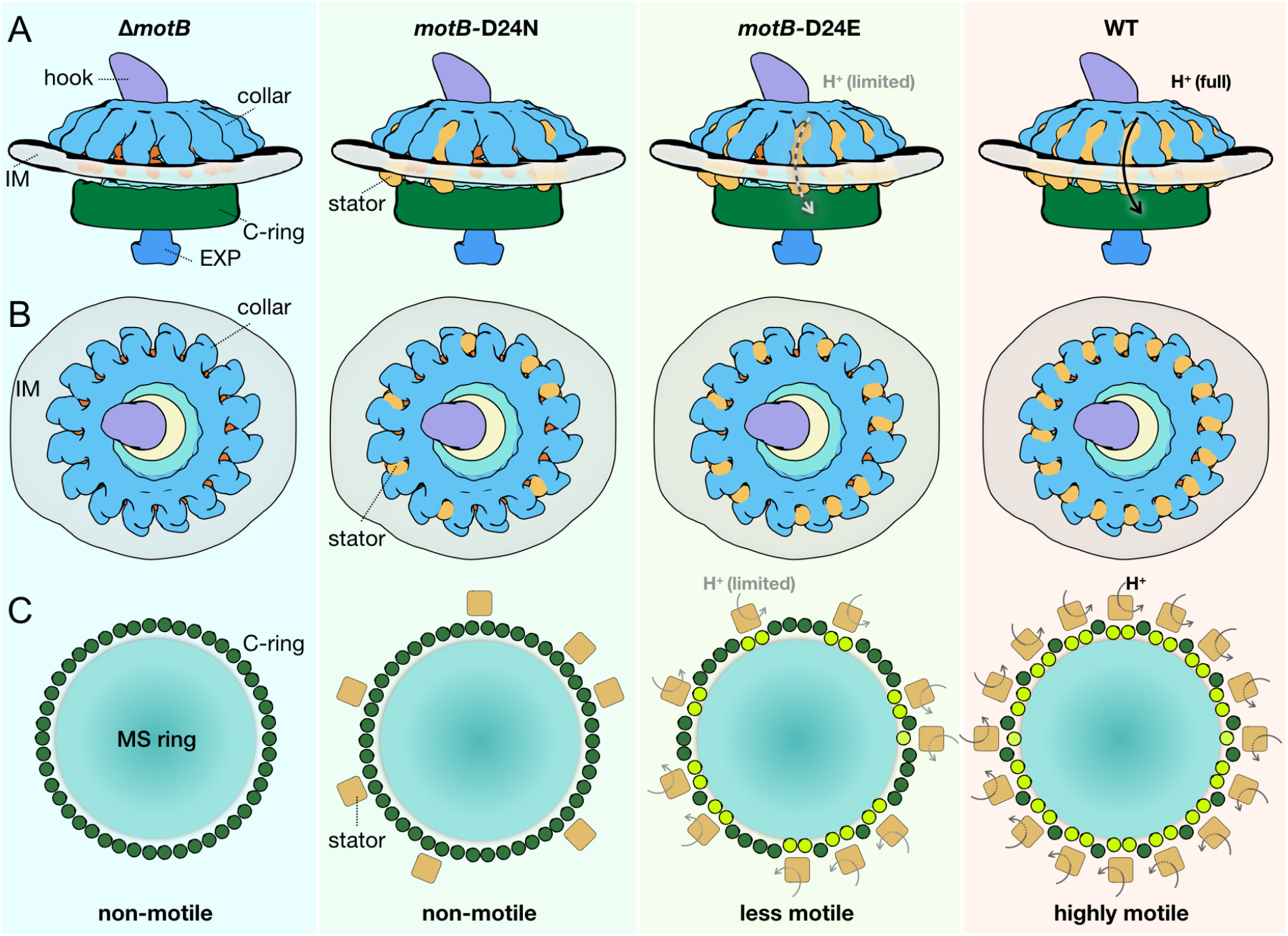
Schematic of stator assembly and stator-rotor interactions. (A, B) Side and top views of stator assembly. A flagellar motor without stators is shown in the left panels. The WT motor with 16 fully assembled stators is shown in the right panels. Several key flagellar components are annotated: C-ring (green), export apparatus (EXP), collar (light blue) embedded in the inner membrane (IM), and hook. Two intermediates in stator assembly show partial occupancy of motors with blocked or attenuated proton conduction by the torque-generating units. (C) Top views of the C-rings from the motors shown in panels A and B. Without proton conduction (as shown in the *motB*-D24N mutant), some stators (tan squares) interact with the FliG units (colored in dark green circles), yet there is relatively little conformational change in the C-ring. When protons flow through the stator channels, the torque generated by the stator induces conformational changes in the FliG units with which they are in contact (colored in light green). In the *motB*-D24E motor, fewer stator units are engaged, and they have a decreased proton flow. As a consequence, the deformations of the C-ring are not as large as in the WT motor, in which the 16 stator units assembled around the C-ring are rapidly conducting protons and generating torque. We propose that the increasing deformation of the C-ring observed with increasing number and activity of assembled stators reflects conformational changes induced by the power strokes of the cytoplasmic MotA loops pushing against the FliGc domain.

In summary, high resolution *in situ* structural analysis of the flagellar motor from wild-type and mutant *B. burgdorferi* provided new insights into the assembly of torque-generating units into the motor and their interactions with other flagellar components in both the periplasm and cytoplasm. The collar structure, which is a unique component of the flagellar motor of spirochetes, acts with the transmembrane protein FliL to help recruit and stabilize sixteen stator complexes. The interactions between the stator complexes and the C-ring result in a large conformational change in the C-ring probably reflects the interaction of the cytoplasmic loop of MotA with FliG_C_ in the proton-flow-driven power stroke that drive rotation of the C-ring.

## EXPERIMENTAL PROCEDURES

### Bacterial strain and growth conditions

High-passage, avirulent *B. burgdorferi sensu stricto* strain B31A and its isogenic mutants (Table S1) were grown in Barbour-Stoenner-Kelly medium without gelatin (BSK II) or plating BSK medium containing 0.5% agarose at 35°C in a 2.5% CO_2_ humidified incubator.

### Construction of Δ***motA*, Δ*motB, motB*D24E*, and motB*D24N and complementation of the Δ*motB* mutant**

Constructions of **Δ***motB* (gene locus number *bb0280*) and **Δ***motA* (locus number *bb0281*) were described previously ^*47*^. Shuttle vector pBSV2G, which carries the gentamicin cassette, was used to complement the *motB::aadA* mutant (Δ*motB*) using native *motB* promoter, *flgB*. The *flgB* promoter and *motB* gene sequences were amplified with primers containing restriction enzyme sites *Xba*I and *Nde*I (5’-3’ and 3’-5’ respectively) and inserted into the *Nde*I site (the 3’ end of the promoter fragment and the 5’ end of the *motB* gene) to yield pFlgBMotB. The primers used were (5’-3’): for *flgB*, FlgB-XbaI-F: <u>tctaga</u>gccggctaatacccgagc and FlgB-NdeI-R: <u>catatg</u>gaaacctccctcatttaa; and for *motB*, MotB.com-F: <u>catatg</u>gctttgcgaattaaga and MotB.com-R: <u>tctaga</u>ttactgcttaatttcctt. Underlined sequences indicate restriction sites. The *flgBmotB* DNA was then ligated into the *Xba*I site of pBSV2G to yield pMotB.com. Two point mutations were generated in MotB (aspartate to glutamate and asparagines, respectively) were constructed. We used pMotB.com plasmid as the PCR template for these substitutions using a site-directed mutagenesis kit (QuikChange, Stratagene Inc.) yielding plasmids MotB(D24E) and MotB(D24N), respectively. These plasmids were sequenced to verify the substitutions. PCR primer sequences for point mutations are given below (5’-3’), and underlined sequences indicate point mutated sequences:

P11 – gttgacttatgga<u>gaa</u>atggttactttgctg

P12 – cagcaaagtaaccat<u>ttc</u>tccataagtcaac

P13 – gttgacttatgga<u>aat</u>atggttactttgctg

P14 – cagcaaagtaaccat<u>att</u>tccataagtcaac.

Approximately 20 µg of purified pMotB.com, MotBD24E, and MotBD24N plasmids were used to transform competent Δ*motB* by electroporation as described above. Transformants were selected with 40 µg/ml gentamicin plus 80 µg/ml streptomycin. Gentamicin and streptomycin resistant clones were confirmed by PCR as well as by western blotting to determine the restoration of MotB synthesis.

### Gel electrophoresis and western blot analysis

Sodium dodecyl sulfate-polyacrylamide gel electrophoresis (SDS-PAGE) and western blotting with an enhanced chemiluminescent detection method were carried out as described previously (45).

### Dark-field microscopy and swarm plate motility assays

*B. burgdorferi* cells (5 × 10^7^ spirochetes/ml) were observed under a dark-field microscope (Zeiss Axio Imager M1), and images were captured using an AxioCam digital camera. Swarm plate motility assays were performed as described ^*47*^. Approximately 1 × 10^6^ cells in a 5 µl volume were spotted onto 0.35% agarose plate containing plating BSK medium diluted 1:10 in Dulbecco’s phosphate buffered saline. Since *B. burgdorferi* is a slow growing organism (8-12 hours generation time), plates were incubated for 5 days at 35°C in a 2.5% CO_2_ humidified incubator. To determine cell morphology, growing *B. burgdorferi* cells were observed under a dark-field microscope (Zeiss Axio Imager. M1). Dark-field images and cells real-time videos were captured using an AxioCam digital camera connected to the microscope. Exponentially growing cells were mixed with 0.5% 400 mesh methylcellulose (Sigma-Aldrich Co.) and video recorded at room temperature (23°C).

### Cryo-EM sample preparation

Cultured WT and mutant cells were centrifuged in 1.5 ml tubes at 5,000 × g for 5 minutes and the resulting pellet was rinsed gently with PBS, then, suspended in 40 µl PBS at a final concentration ∼2×10^9^ cells/ml ^*33*^. After mixing with 10 nm gold fiducial markers, 5 µl *B. burgdorferi* samples were deposited onto freshly glow-discharged holey carbon grids. Grids were blotted with filter paper and then rapidly frozen in liquid ethane, using a home-made gravity-driven plunger apparatus.

### Cryo-electron tomography

The frozen-hydrated specimens were transferred to a 300-kV Polara G2 electron microscope (FEI) equipped with a Direct Electron Detector (DDD) (Gatan K2 Summit) or with a charge-coupled-device (CCD) camera (TVIPS; GMBH, Germany). Images were recorded at 15,400× magnification with pixel size of 2.5 Å (for images recorded by K2) or at 31,000× magnification with pixel size of 5.7 Å (for images recorded by CCD). SerialEM ^*57*^ was used to collect tilt series at −6 to −8 µm defocus, with a cumulative dose of ∼100 e^-^/Å^2^ distributed over 61 images and covering angles from −60°to 60°, with a tilt step of 2°. Images recorded by K2 camera were first drift-corrected using the *motioncorr* program ^*58*^. Then all tilt series were aligned using fiducial markers and volumes were reconstructed by the weighted back-projection method using IMOD ^*59*^.

### Subtomogram analysis

Bacterial flagellar motors were manually picked from the tomograms as described ^*60*^. The subtomograms of flagellar motors were extracted from the tomograms. In total, 14,049 subtomograms were manually selected from the tomographic reconstructions and used for subtomogram analysis. The i3 software package ^*61, 62*^ was used to generate the average motor structure. Class averages were computed in Fourier space so that the missing wedge problem of tomography was minimized ^*62, 63*^. Fourier shell correlation coefficients were calculated by generating the correlation between two randomly divided halves of the aligned images used to generate the final maps.

For the local alignment around the stator region, regions around the sixteen stators in each motor were first extracted and aligned, then classified based on the stator, the C-ring and cytoplasmic membrane features. Only those containing an almost flat region of the cytoplasmic membrane were selected for further refinement. Number of subtomograms used for local alignment were listed in Table S2.

### Measurement of the C-ring tilt angles in the Δ***motB, motB*-D24N, *motB*-D24E and WT motors**

To measure the C-ring tilt angles in average, 16-fold symmetry was first applied to the Δ*motB, motB*-D24N, *motB*-D24E and WT motor structures. We selected the cross sections of the motor structures. Treat the C-ring density (without FliG_N_ density) in the cross-section images as a whole object, then calculate an ellipse that can fit the object shape. The angle between the long axis of the ellipse and the Y-axis was considered as the tilt angle of the C-ring.

### Three-dimensional visualization and modeling

UCSF Chimera ^*64*^ and ChimeraX ^*65*^ software packages were used for surface rendering of subtomogram averages and molecular modeling. For the surface rending of WT motor structure, all stator densities are from the local alignment results shown in Figure 3I, then fitted into the motor density shown in Figure 2D through the function “fitmap” in Chimera or ChimeraX, thus the 16 stators are almost the same, just have relatively different orientations and positions. For the surface rendering of Δ*motB* motor structure, the density map shown in Figure 2A was used. The crystal structures of FliG_N_ (PDB ID: 5TDY), FliM_C_-FliG_MC_ complex (PDB ID: 3HJL) and FliN ^*52*^ were docked into the density map through the function “fitmap”.

## ACKNOWLEDGEMENTS

We thank Mike Manson for critical reading and suggestions. This research was supported by grants R01AI087946 (to J.L.) and R01AI132818 (to M.M.) from the National Institute of Allergy and Infectious Diseases and R01GM107629 from the National Institute of General Medicine (to J. L.).

## SUPPORTING INFORMATION

**Figure 1-figure supplement 1.**
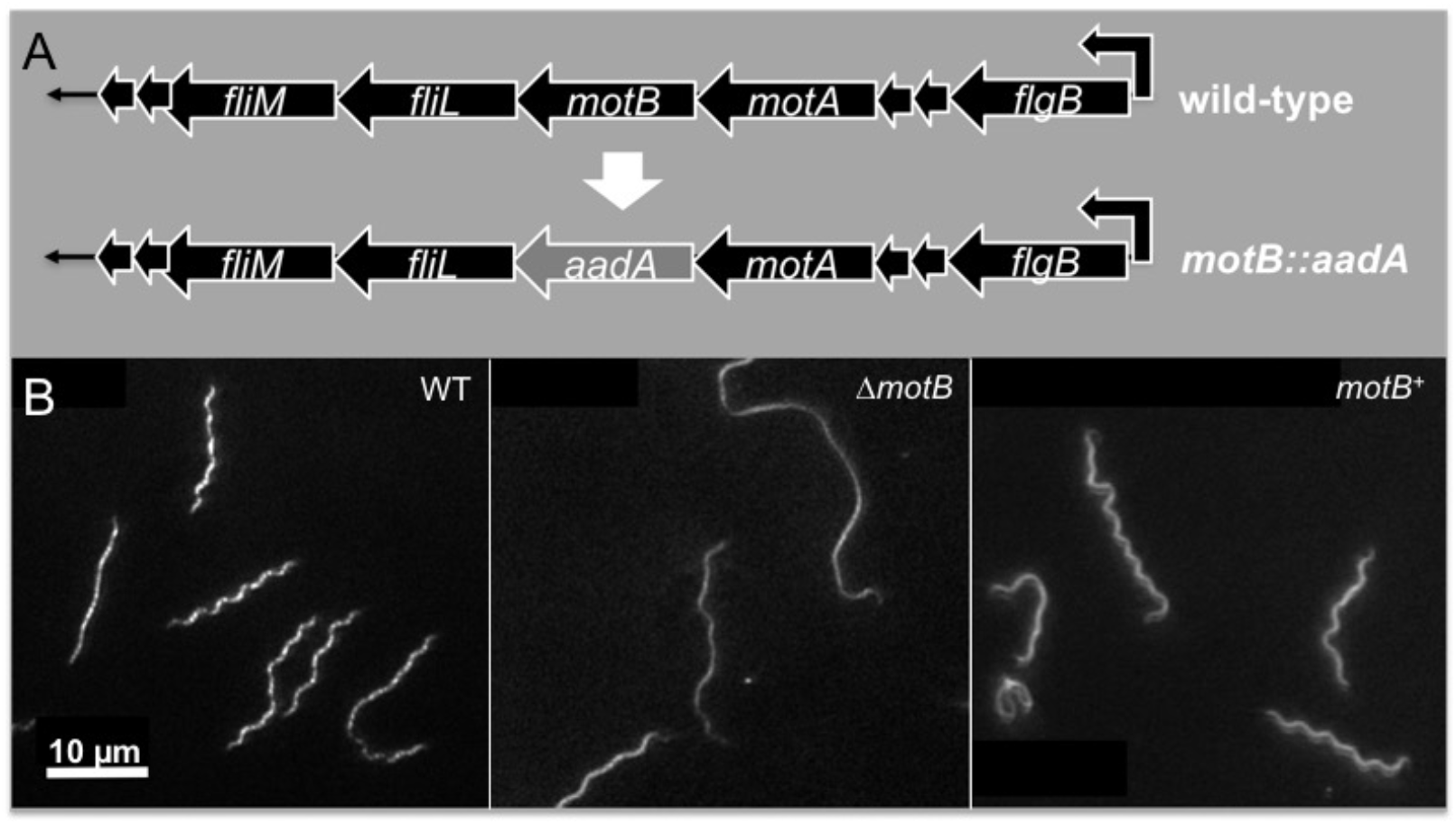
The motilities and cell morphologies of WT, Δ*motB* and *motB*^+^ *B. burgdorferi* cells.

**Figure 1-figure supplement 2.**
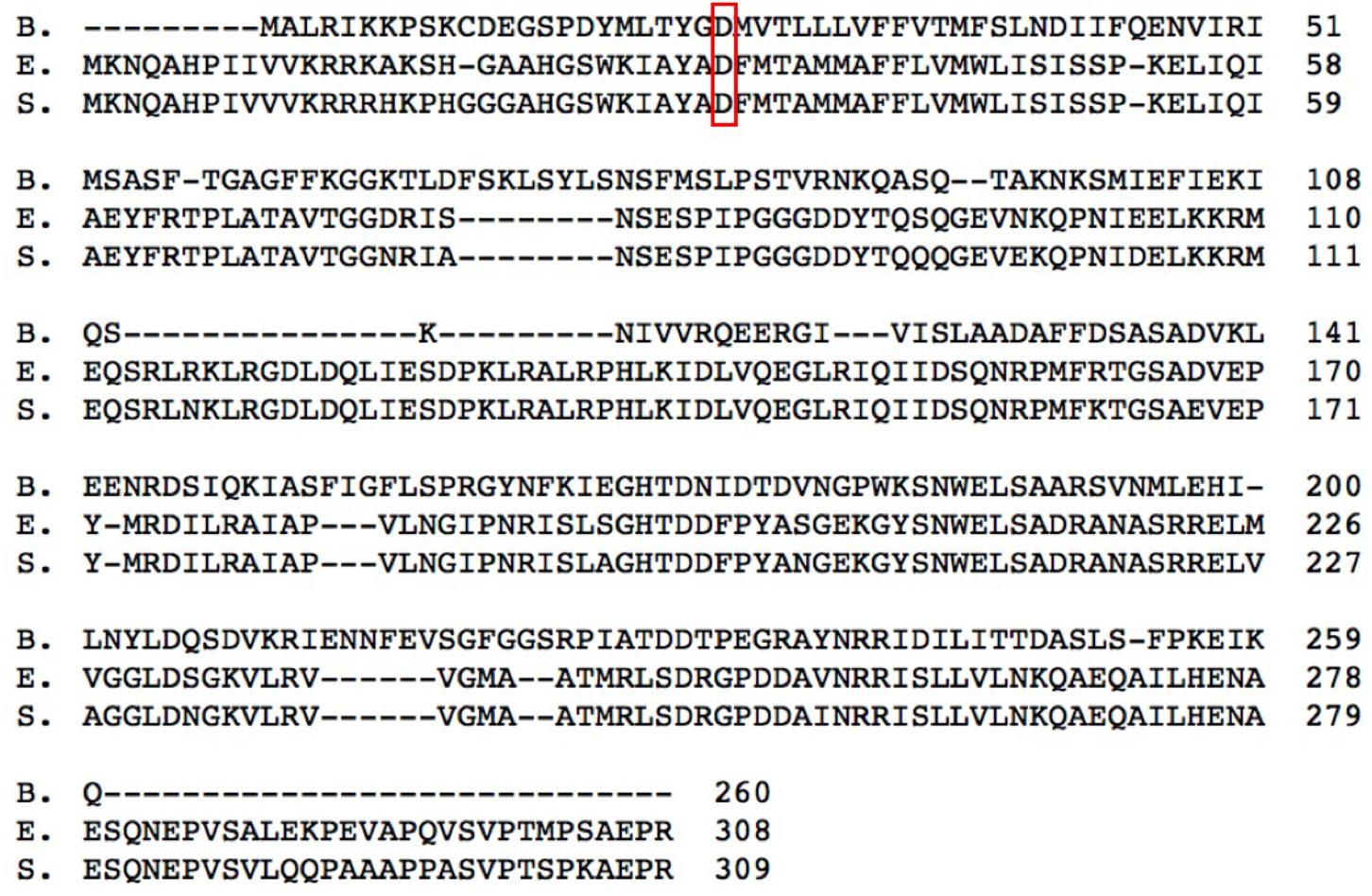
Sequence alignment of *motB* genes in *Borrelia burgdorferi* (B.), *Escherichia coli* (E.) and *Salmonella enterica* (S.). The conserved aspartic acid residue D (24 in B., 32 in E. and 33 in S.) is indicated by red frame.

**Figure 2-figure supplement 1.**
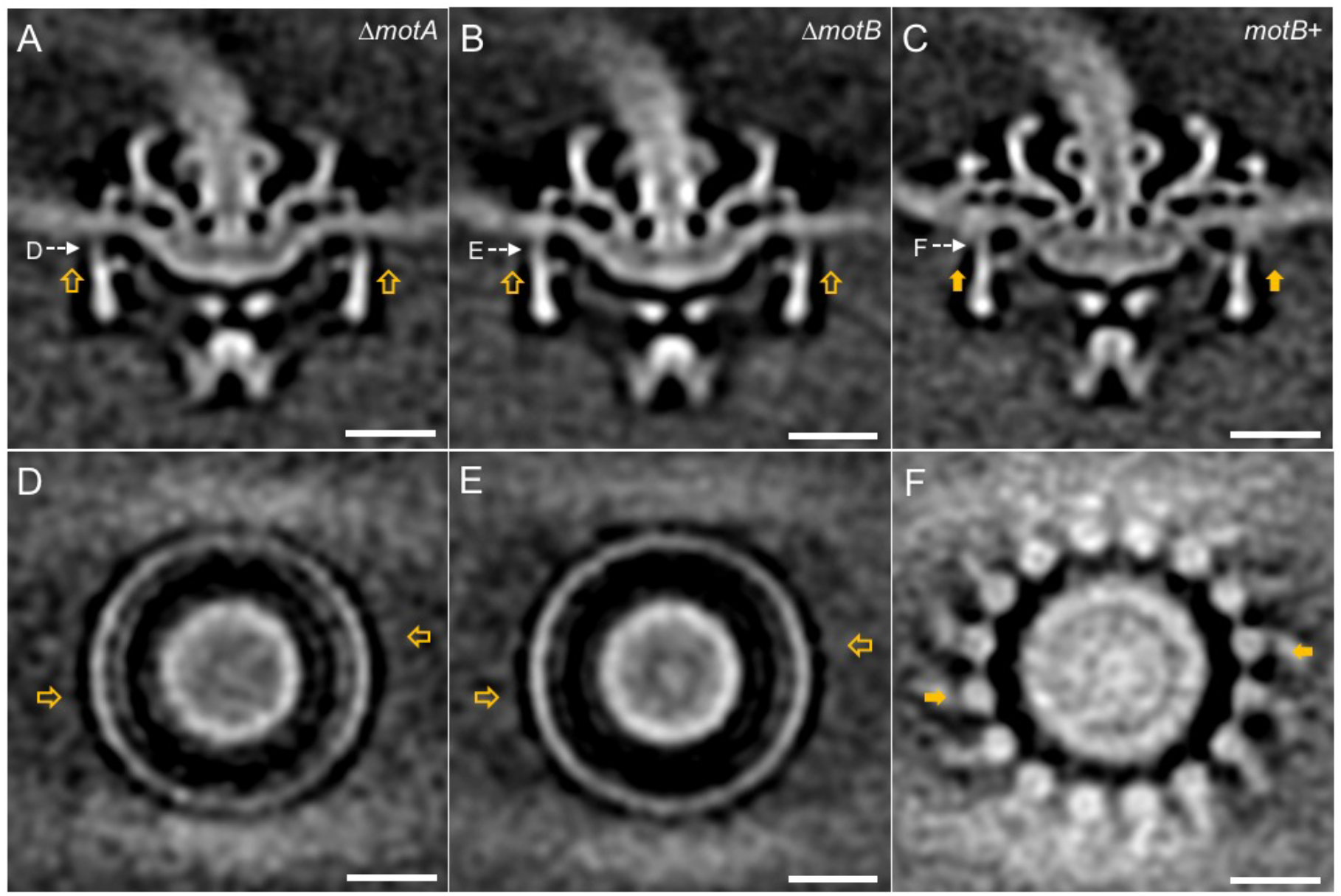
Asymmetric reconstructions of Δ*motA*, Δ*motB* and *motB*^+^ motors in *B. burgdorferi*. (A-C) A central section of the asymmetric reconstruction of the Δ*motA*, Δ*motB* and *motB*^+^ motor structure, respectively. (D-E) A cross section from the asymmetric reconstruction of Δ*motA*, Δ*motB* and *motB*^+^ motor structure at the top of the C ring, respectively. Two stators in *motB*^+^ motor structure are indicated by solid arrows, and the corresponding positions in Δ*motA* and Δ*motB* are indicated by empty arrows. Bar = 20 nm.

**Figure 3-figure supplement 1.**
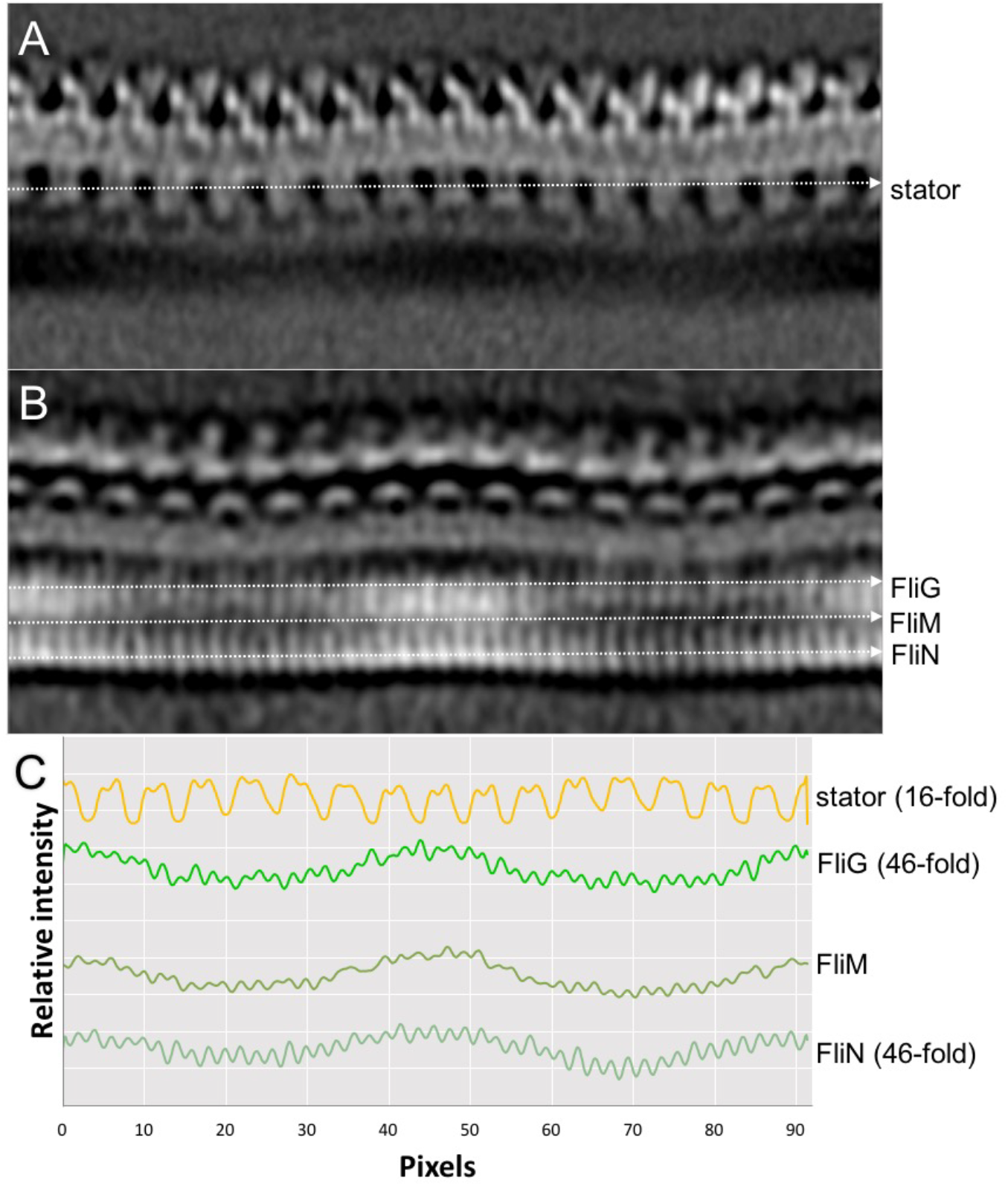
Symmetries of the stator and the C-ring in WT motor structure. (A, B) Two sections of the map of an unrolled WT motor structure (unroll along its central rod). (C) Intensity plot files along the stator, FliG, FliM and FliN densities (indicated by dashed lines in A and B) showing their symmetries.

**Figure 4-figure supplement 1.**
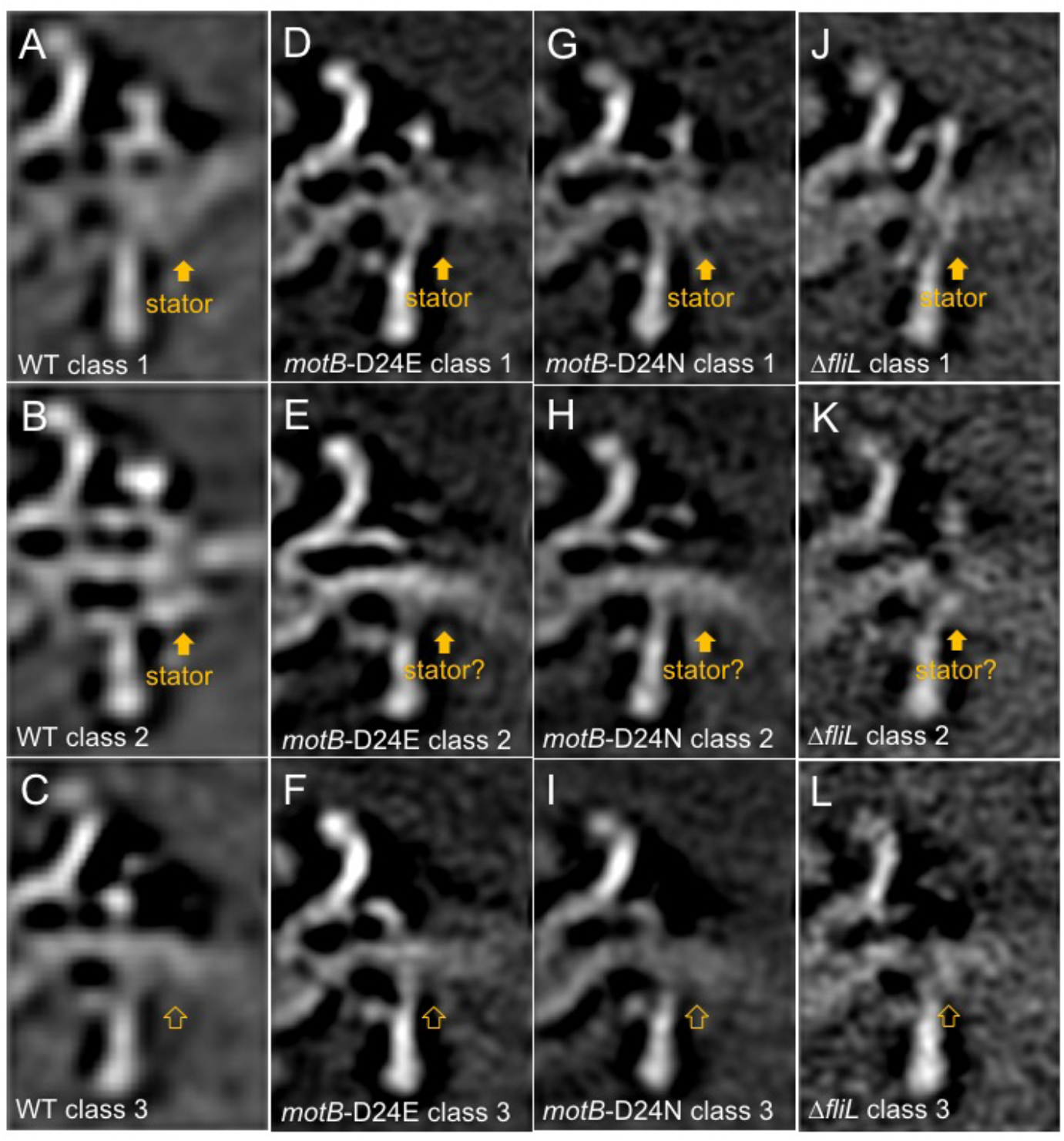
Measurement of stator occupancies in WT, *motB*-D24E, *motB*-D24N and Δ*fliL* motors. 3D classification was applied to WT, *motB*-D24E, *motB*-D24N and Δ*fliL* motor structures based on the stator density, then the stator occupancy was calculated by dividing the particle numbers in class averages with stator densities over the total particle numbers. Three selected class averages from the classification results of WT (A-C), *motB*-D24E (D-F), *motB*-D24N (G-I) and Δ*fliL* (J-L) motor structures are shown here. (A, B, D, G and J) show class averages with stators. (C, F, I and L) show class averages without stators. (E, H and K) show class averages that we are not certain whether there are stators or not.

**Figure 4-figure supplement 2.**
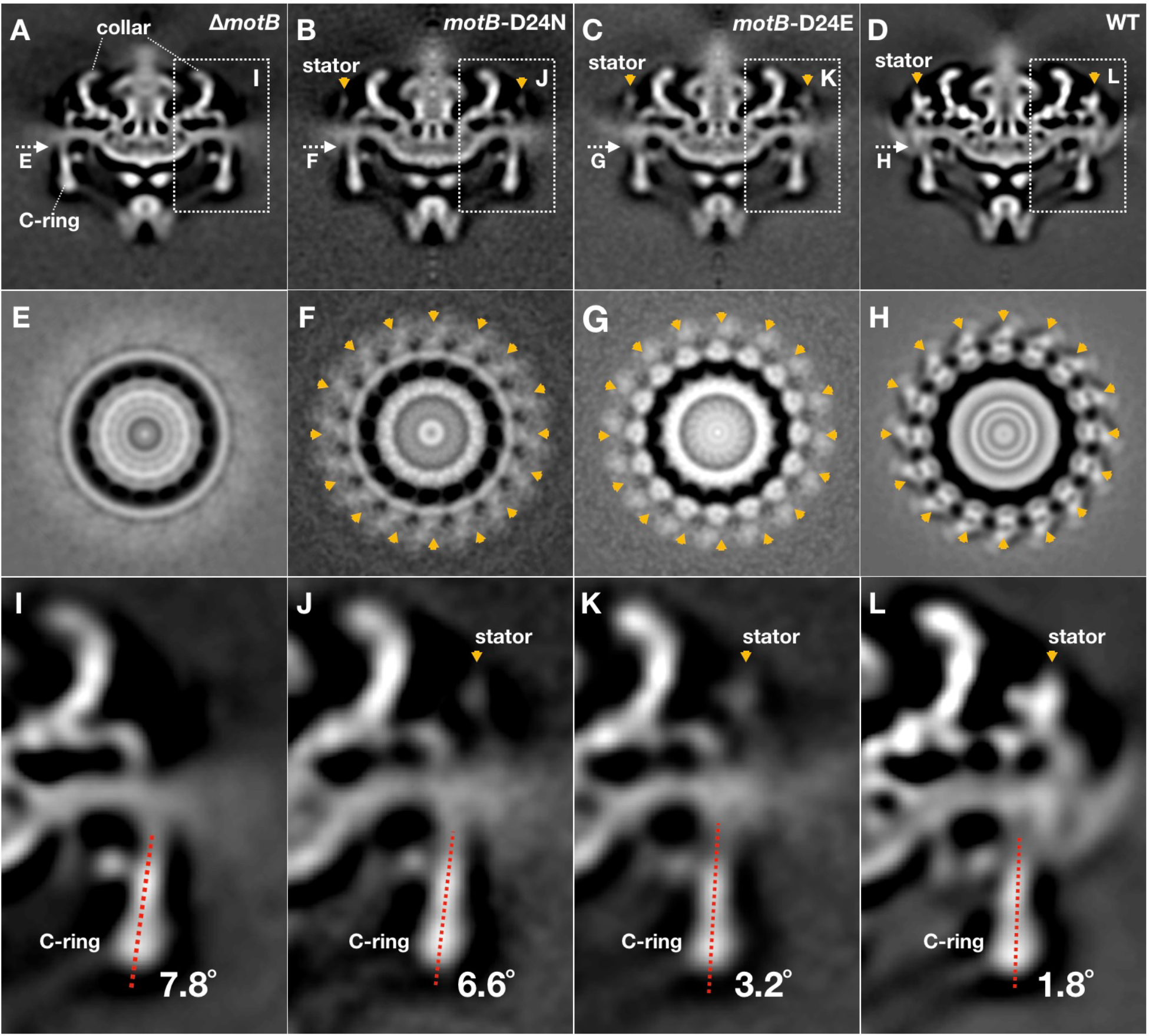
Conformational changes of the C-ring induced by stator-binding and proton flux. (A-D) A central section of Δ*motB, motB*-D24N, *motB*-D24E and WT motor structures (16-fold symmetry was applied), respectively. Stators are indicated by orange arrows. (E-F) A cross-section at the top of the C-ring from the Δ*motB, motB*-D24N, *motB*-D24E and WT motor structures, respectively. Stators are indicated by orange arrows. The C-ring undergoes considerable modulation at the stator binding positions in *motB*-D24E (G) and WT (H) motors. (I-L) The tilt angles of the C-ring away from the rotation axis in the Δ*motB, motB*-D24N, *motB*-D24E and WT motor structures, respectively.

**Figure 4-figure supplement 3.**
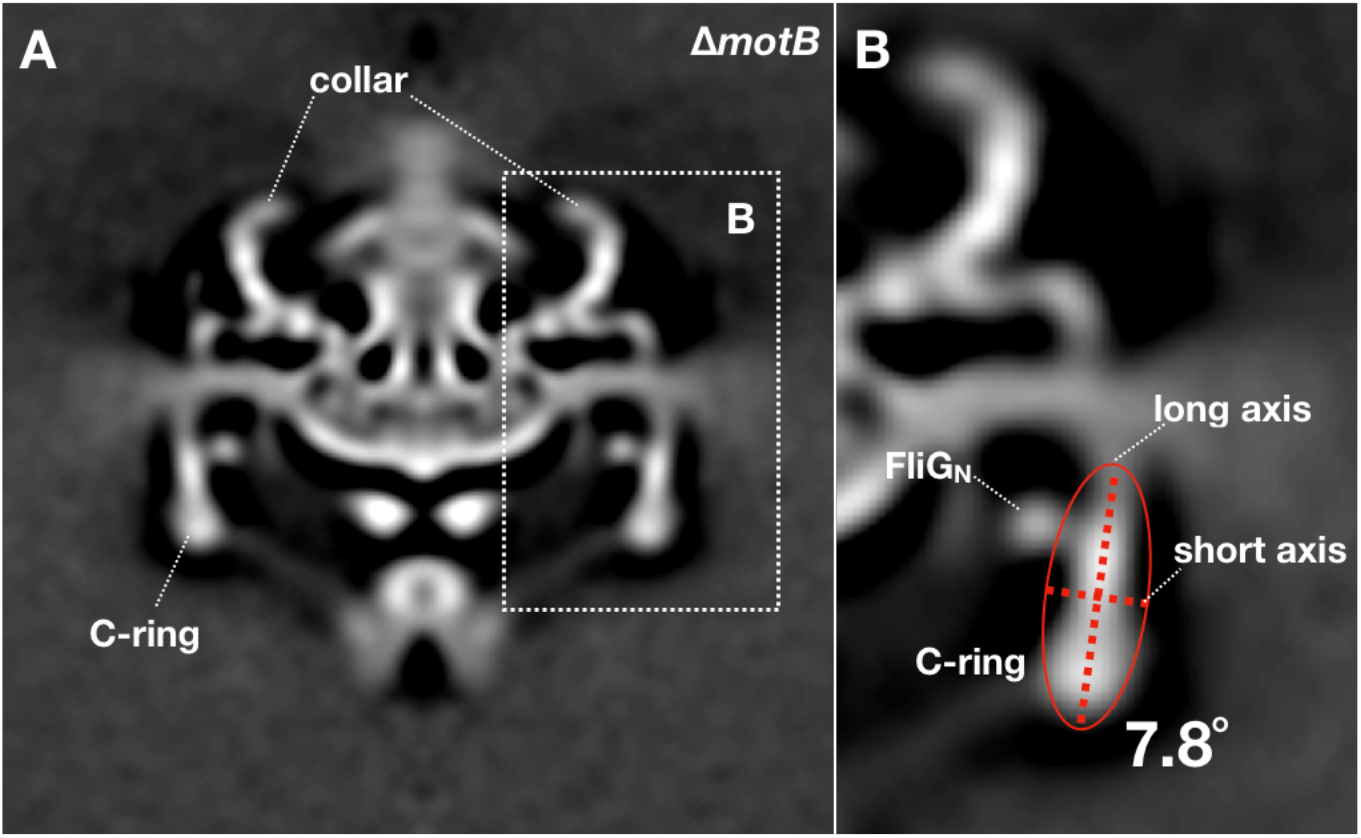
Measurement of the C-ring tilt angle in the Δ*motB* motor structure. 16-fold symmetry was applied to the Δ*motB* motor structure. (A) A cross section of the symmetrized Δ*motB* motor structure. (B) Treat the C-ring density (ignore FliG_N_ density) as a whole object, then calculate an ellipse that can fit the object shape. The angle between the long axis of the ellipse and the Y-axis was considered as the tilt angle of the C-ring (7.8°). The tilt angles of the C-ring in *motB*-D24N, *motB*-D24E and WT motor structures were measured by the same method.

**Figure 4-figure supplement 4.**
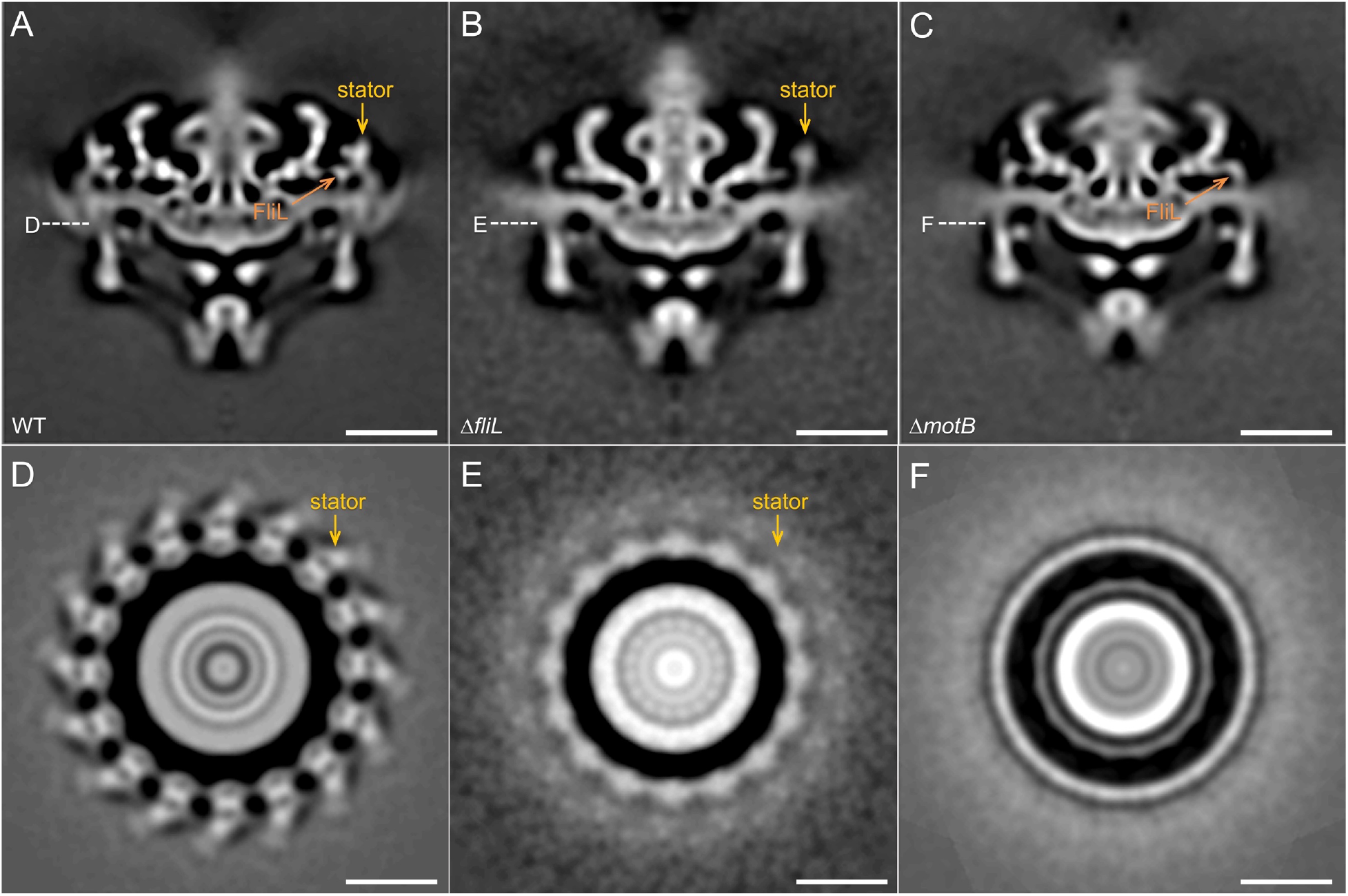
Comparison of the motor structures from WT, Δ*fliL* and Δ*motB*. (A-C) A central section of the WT, Δ*fliL* and and Δ*motB* motor structures (16-fold symmetry was applied), respectively. Stators are indicated by orange arrows. (D-F) A cross-section at the top of the C-ring from the WT, Δ*fliL* and and Δ*motB* motor structures, respectively. Stators are indicated by orange arrows. Note that the stator densities in the Δ*fliL* motor is relatively weak compared to those in the WT motor, suggesting that there are fewer stator units in the Δ*fliL* motor.

**Table S1.** Strains used in this study.

| Strains | Gene | Gene product | Cell morphology | PFs morphology | Motility | Reference |
| --- | --- | --- | --- | --- | --- | --- |
| Wild type | N/A | N/A | spiral shaped | flat ribbon | motile | Liu. *et al.* (2009) |
| $\Delta motB$ | BB0280 | stator protein | rod shaped | flat ribbon + disrupt ribbon | nonmotile | This study |
| $\Delta motA$ | BB0281 | stator protein | rod shaped | flat ribbon + disrupt ribbon | nonmotile | This study |
| *motB*-D24E | N/A | N/A | spiral shaped | flat ribbon | less motile | This study |
| *motB*-D24N | N/A | N/A | rod shaped | flat ribbon | nonmotile | This study |
| *MotB*<sup>+</sup> | N/A | N/A | spiral shaped | flat ribbon | motile | This study |
| $\DeltafliL$ | BB0279 | motor protein | spiral shaped | flat ribbon | less motile | Motaleb *et. al.* (2011) |

**Table S2.**
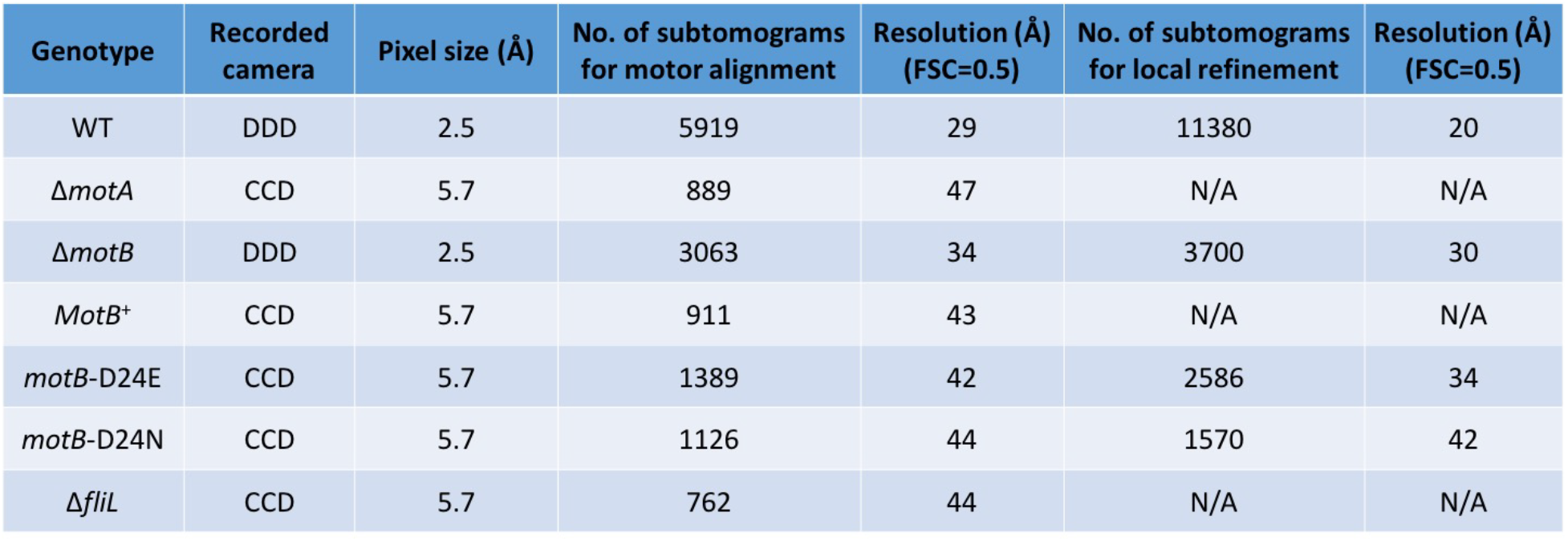
Cryo-ET data, parameters used in this study and the resolution of subtomogram average structures.

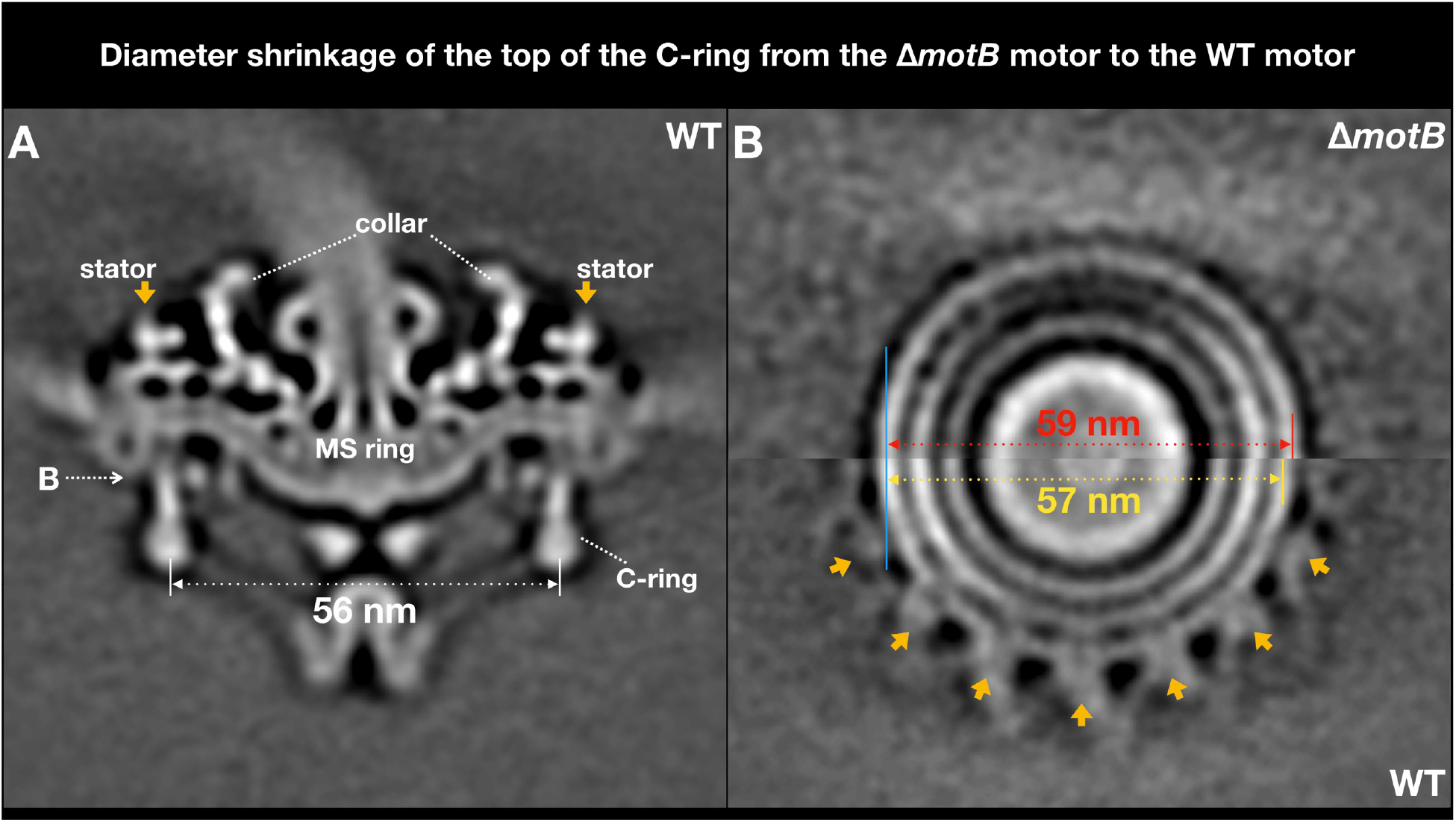

**Video 1. Diameter shrinkage of the top of the C-ring from the Δ*motB* motor to the WT motor.**

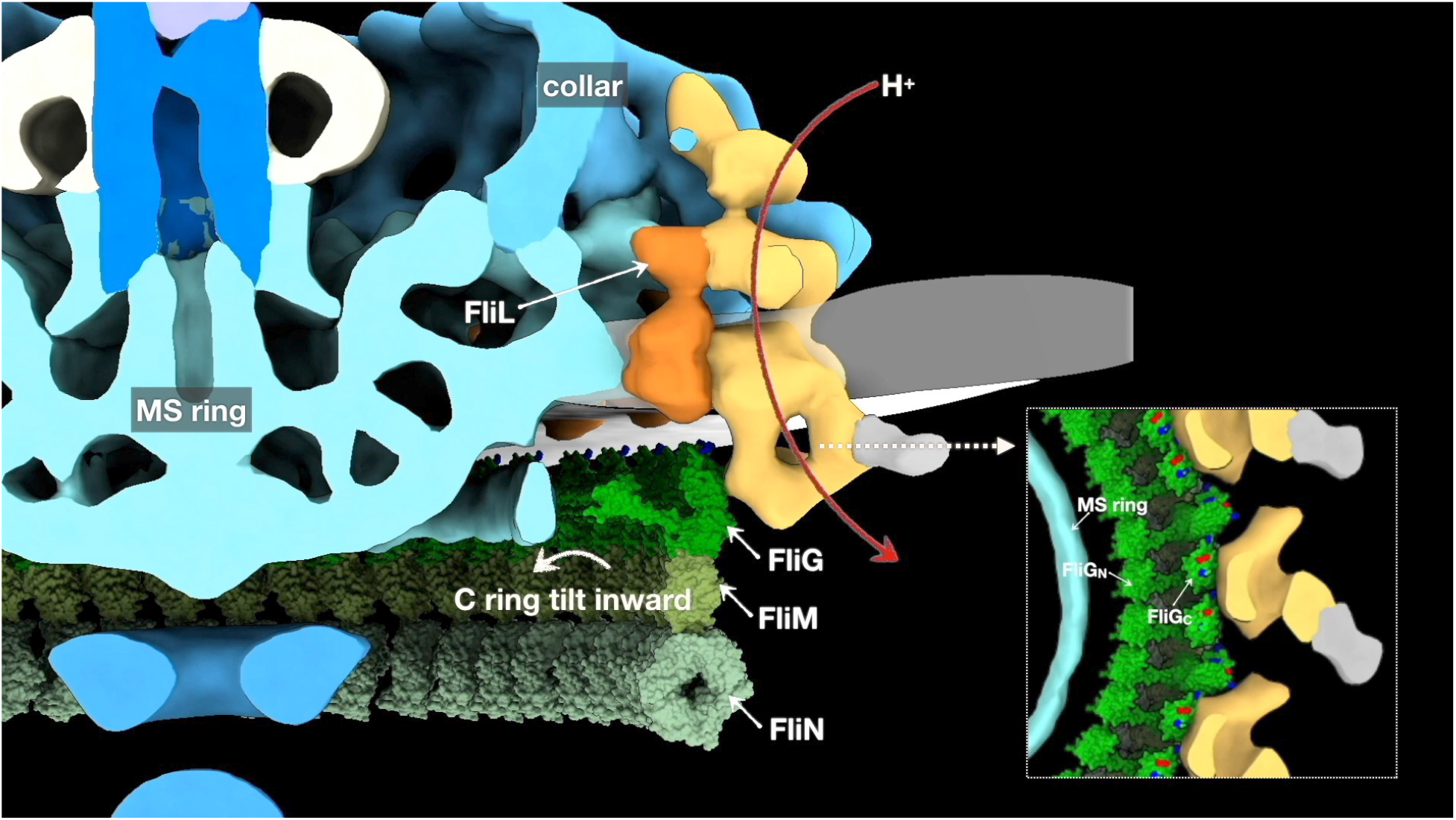

**Video 2. *In situ* structural analysis of the stator-rotor interactions in *Borrelia*.**

